# Infant functional connectivity fingerprints predict long-term language and pre-literacy outcomes

**DOI:** 10.1101/2020.10.29.360081

**Authors:** Xi Yu, Silvina Ferradal, Danielle D. Sliva, Jade Dunstan, Clarisa Carruthers, Joseph Sanfilippo, Jennifer Zuk, Lilla Zöllei, Emma Boyd, Borjan Gagoski, P. Ellen Grant, Nadine Gaab

## Abstract

Functional brain networks undergo extensive development within the first few years of life. Previous studies have linked infant functional connectivity to cognitive abilities in toddlerhood. However, little is known regarding the long-term relevance of functional connections established in infancy for the protracted development of higher-order abilities of language and literacy. Employing a five-year longitudinal imaging project starting in infancy, this study utilizes resting-state functional MRI to demonstrate prospective associations between infant functional connectivity fingerprints and subsequent language and foundational literacy skills at a mean age of 6.5. These longitudinal associations are preserved when key environmental influences are controlled for and are independent of emergent language abilities in infancy, suggesting early development of functional network characteristics in supporting the acquisition of high-order language and pre-literacy skills. Altogether, the current results highlight the importance of functional organization established in infancy as a neural scaffold underlying the learning process of complex cognitive functions.

## Introduction

Human brains undergo extensive changes within the first years of life, establishing critical foundations of neuroarchitecture that underlies the protracted maturational process of perceptual and higher-order cognitive functions. Recent advances in pediatric neuroimaging techniques refine functional mapping of the infant brain, primarily through resting-state functional connectivity (FC) MRI analyses [1]. FC characterizes functional brain organization by examining synchronized fluctuations in the blood oxygen dependent (BOLD) signals among different brain regions [2]. For instance, FC analyses conducted in late pre-term and full-term neonates have reported adult-like functional connectivity patterns in regions underlying motor and primary sensory processing (e.g., primary auditory and visual networks) at birth [3–6]. As maturation progresses, long-distance functional connectivity patterns among association cortices develop, coupled with the emergence of higher-order cognitive functions such as executive function and language [5–8]. To date, only a few studies have directly examined the cognitive developmental significance of functional connectivity emerging in infancy, revealing associations between infant FC patterns and cognitive abilities and pathophysiology in toddlers and preschoolers [9–11]. However, it is still unknown whether and how FC in infancy can account for individual differences in the long-term trajectory of language and foundational literacy abilities into school-age.

Early language acquisition starts in utero and is significantly shaped by the linguistic environment [12]. Newborns demonstrate selective preferences for human speech compared to other sounds, such as sine-wave analogs with comparable spectral and temporal parameters as those of real speech (e.g., [13, 14]), and are able to recognize and categorize different aspects of speech, such as phonetic and stress patterns [15–17]. Infants typically develop an understanding of familiar words as early as the second half of the first year, building a critical foundation for long-term language development into school-age [18–21]. Early linguistic experiences further serve as critical building blocks for the development of emergent literacy skills. These skills include phonological processing, the ability to recognize and manipulate the internal structure of speech sounds, one of the key predictors of subsequent reading development (e.g., [22–24]). Moreover, this developmental time course is sensitive to environmental characteristics, including language and early literacy exposure. For example, it has been shown that the quality and quantity of child-directed speech are highly correlated with vocabulary gain in toddlers [25–27]. In addition to language exposure, home literacy practices are also associated with subsequent verbal and literacy skills during preschool and school-age (e.g., [28–31]), highlighting the important role of the environment. Moreover, emergent literacy skills are then further deepened through instruction once the child starts school and reciprocally influence language development, such as vocabulary [32–35]. Overall, the development of language and pre-literacy skills starts in infancy and continues through childhood and beyond, constantly shaped by the quantity and quality of input provided in the child’s home, school, and other environmental contexts.

Crucially, at the time of birth, infant brains have already developed the essential neurobiological scaffold for language and literacy acquisition. It has been repeatedly demonstrated that, when listening to human speech compared to control sounds or speech played backward, newborns and preverbal babies activate language-related areas [36, 37], largely located in the temporal lobe [38–41]. Electrophysiological responses to speech perception in infancy are prospectively associated with language and literacy outcomes and trajectories at preschool and school-age, suggesting early brain function has a long-lasting influence on subsequent language and literacy acquisition (e.g., [42–48]). Moreover, white matter characteristics have been shown to correlate with the concurrent verbal skills in the first five years of life [49], and white matter organization at term age has been shown to predict preterm infants’ language development at age two [50], implying the early emergence of structural connectivity underlying language development. Furthermore, it has been shown that ***the developmental trajectories of large-scale functional topologies*** within the first two years of life are associated with overall developmental progress on language milestones (i.e., functional-equivalent age) at age 4 in a group of 23 children [51]. These results demonstrate the importance of early changes in functional topologies on language development. However, it is still an open question as to what extent functional network characteristics established in early in infancy are prospectively associated with individual differences in different aspects of language and literacy abilities, such as vocabulary and phonological processing, after children receive formal instruction at school age. Moreover, it is unknown whether such long-term associations, if observed, are driven by (pre)language skills and early linguistic environmental variables, or reflect early brain mechanisms specifically linked to the high-order cognitive function of language and literacy skills.

Employing a five-year longitudinal imaging dataset from infancy to school-age, the current study utilizes resting-state functional connectivity MRI methods to investigate prospective associations between infant functional brain networks and subsequent measures of language and emergent literacy skills at age 6. We first conducted multivariate pattern analysis (MVPA) to identify functional networks in infancy that are associated with subsequent school-age performance while controlling for critical environmental factors. Further analyses were performed to assess whether the inclusion of receptive and expressive language performance measured in infancy leads to changes in these observed associations. The results of the current study offer the potential to inform our understanding of how early-emerging functional network mechanisms can contribute to individual differences in language and reading abilities later in life.

## Results

### Infant FCFs can predict school-age language and foundational literacy skills

We first evaluated whether functional connectivity fingerprints (FCF) in infancy were prospectively associated with the subsequent development of language and foundational literacy skills at school-age. The outcome measures included oral language (OL) and two key pre-literacy skills commonly recognized as crucial cognitive precursors of reading – phonological processing and rapid automatized naming (RAN, [52–54]). To probe the functional networks specifically associated with long-term language and literacy development, special consideration was taken to minimize the impact of other factors known to influence children’s verbal skills. Specifically, key environmental factors, including socio-economic status (SES; indicated through parental educational background) and home literacy environment (HLE), were collected during the infant imaging time point (see SI Methods) and included as covariates along with the infant age at the scan. We further measured the children’s general cognitive abilities subsequently at school-age, which were included as a control variable to partial out individual variance in general cognitive maturation. Meanwhile, given the close associations between the resting-state FC pattern of one region and the functional relevance (i.e., task-based fMRI activation) of this region [55–57], region-specific FC patterns were characterized for each infant brain using the FCF approach that computed whole-brain functional connectivity pattern associated with each anatomical region of interest (ROI).

To identify infant functional networks prospectively associated with school-age performance of language and pre-literacy skills, Support Vector Regression (SVR) analyses were performed for each pair of the region-specific FCFs and each measure of language and pre-literacy skill (see Figure 1 for a full description of the analysis). For every infant, region-specific FCF of each ROI was generated using Pearson’s correlations between BOLD time series of this region and that of all cortical ROIs spanning the entirety of the cortex derived from the one-year-old infant AAL atlas [58]; see a full list of all 78 regions in table S2). The current analysis focused on the FCFs of 16 seed ROIs located in the bilateral temporal lobes, given the important role of temporal lobe regions in speech and language perception tasks in infants [38, 39, 59–61], The FCF associated with a particular seed region was entered into the SVR models as the input feature with performance on each of the three assessments (OL, phonological processing, RAN) as the outcome measures in order to identify infant functional networks important for development in each specific cognitive domain. Moreover, to minimize overfitting, the SVR model was first trained using 80% of the dataset and then tested using the remaining 20% of the data. This process was repeated 1000 times, generating a (true) distribution of the SVR performance. Permutation tests were subsequently performed following the same procedure except that each of the tasks performed was randomly assigned to every participant in order to generate a random/null distribution. Both distributions, obtained by the real and shuffled performance, were compared using a Kolmogorov-Smirnov test, and the effect size was calculated using Cohen’s d [62]. Moreover, a 95% bootstrap confidence interval (BCI) was further estimated for the mean association strength for each FCF-behavioral measure pair based on the true distribution of the SVR performance. Here, we only focus on strong associations with (a) an effect size values of .80 or higher, signaling a large effect based on Cohen (2013), (b) a 95% BCI excluding zero mean, indicating that a null correlation was not plausible, and (c) significantly outperformed the respective randomized distribution, i.e., *p*< 0.05 after correcting for multiple comparisons.

**Figure 1.**
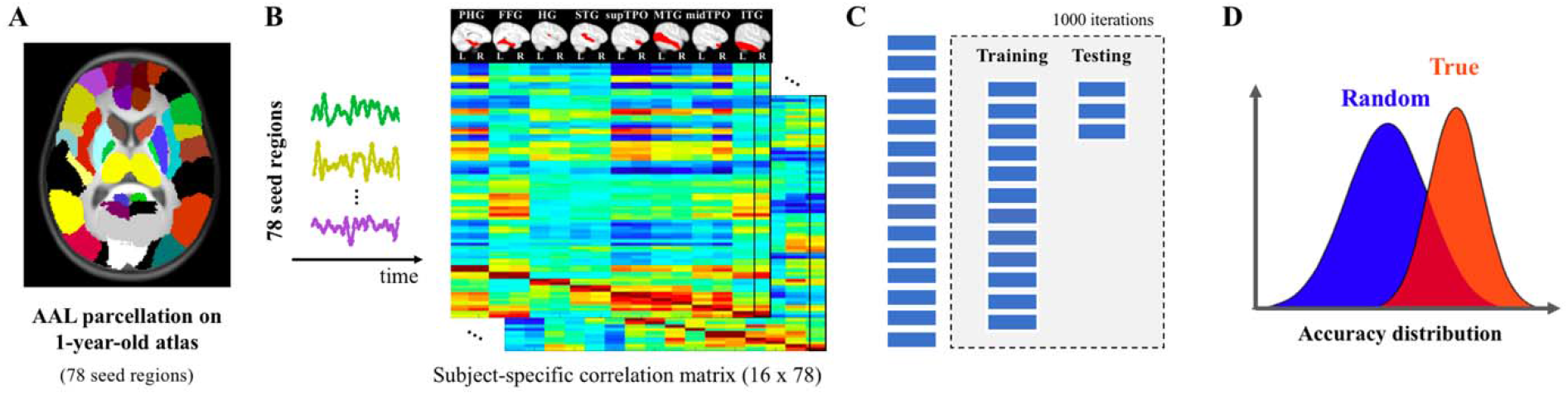
General pipeline of the prediction analyses. **A.** Seed regions were defined as 78 cortical parcellations derived from the 1-year-old infant AAL atlas [58]. **B.** Time series were extracted for every participant by averaging the BOLD signal across all the voxels within each seed region. The functional connectivity fingerprints (FCF) were obtained by pair-wise Pearson’ correlations. The current analyses focused on 16 seed regions residing in the bilateral temporal lobes given the previously reported engagement of this region in speech and language processing in infancy [38, 39, 59–61]. Thus, a 16 × 78 correlation matrix was generated for each subject, where each column represents an FCF pattern associated with a specific seed region in the temporal lobe as shown in the top sagittal views. **C.** Prediction analyses were performed to identify regions whose FCFs in infancy were prospectively associated with the long-term development of language and pre-literacy skills at school-age. Each FCF pattern was entered into a Support Vector Regression (SVR) model as features for the prediction analyses on every school-age performance. To objectively evaluate the prediction performance, the whole sample was randomly divided into training (80%) and testing (20%) datasets. Correction for covariates was first performed in the training dataset only, where the task performance (standard scores) was submitted to a linear regression model with environmental measures (SES and HLE), general cognitive skill at school-age, and age in months at the infant imaging time point as independent regressors. The residuals of the models, representing individual differences for each skill after controlling for environmental and general cognitive influences, were then entered into the SVR analyses. The trained SVR model was then applied to the FCF patterns of the testing datasets, which generated the predicted task performance. The predicted values were then correlated with the true performance (i.e., residual values derived from the linear regression model based on the standard scores of the testing set), and the correlation coefficient was considered as performance of the SVR model. This procedure was repeated 1000 times to produce a distribution of prediction performance for each pair of seed region and task performance. **D.** The distribution of prediction performance based on the true language and literacy outcomes was compared against a null distribution generated by the same procedure, but assigning random scores of school-age performance (permutation tests) using a Kolmogorov-Smirnov test. An effect size and a 95% bootstrap confidence interval were further calculated for the prediction performance of each seed region-behavioral assessment pair. Abbreviations: PHG = ParaHippocampal gyrus; FFG = Fusiform gyrus; HG = Heschl’s gyrus; STG = Superior temporal gyrus; supTPO = Temporal pole (the superior portion); MTG = Middle temporal gyrus; midTPO = Temporal pole (the middle portion); ITG = Inferior temporal gyrus.

The SVR analyses revealed a strong association between the FCF of the middle portion of the left temporal pole (midLTP, *r* = 0.42 ± 0.36, 95% BCI = [0.40, 0.44], *p*_corrected_ < 0.001, Cohen’s d = 1.1) in infancy and oral language skills measured at school-age (Figure 2). We further identified three regions whose FCF patterns exhibited strong associations with phonological processing skills at the school-age: the left fusiform gyrus (LFFG, *r* = 0.40 ± 0.37, 95% BCI = [0.38, 0.43], *p*_corrected_ < 0.001, Cohen’s d = 0.96), right fusiform gyrus (RFFG, *r* = 0.31 ± 0.39, 95% BCI = [0.29, 0.34], *p*_corrected_ < 0.001, Cohen’s d = 0.82), and the right inferior temporal gyrus (RITG, *r* = 0.41 ± 0.36, 95% BCI = [0.39, 0.43], *p*_corrected_ < 0.001, Cohen’s d = 0.94, Figure 2, see the full results in Table S3). Similar result patterns were also obtained based on 30% and 40% test datasets (Table S4), as well as adopting a leave-one-out cross-validation approach (SI Methods and Table S5). No region in the temporal lobe showed significant associations between infant FCFs and subsequent RAN abilities. Lastly, correlation analyses were computed between motion parameters (number of outliers and mean FD) and language/literacy assessments at school-age. No significant results were observed (all *p*_uncorrected_ > 0.1, Table S6), indicating that head movement during image acquisition cannot explain the observed associations.

**Figure 2.**
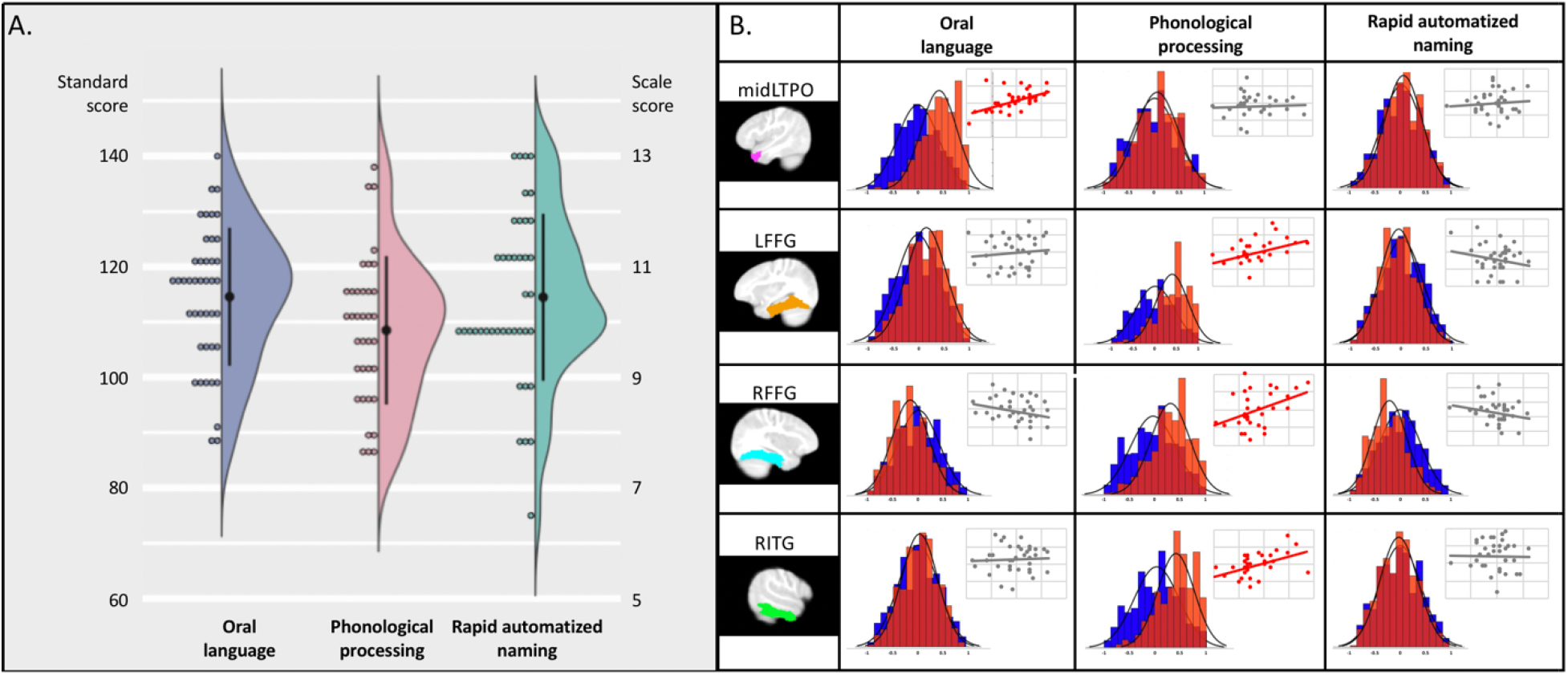
Prospective associations between functional connectivity fingerprints (FCF) in infancy and language and foundational literacy development at school-age. **A.** Distribution of the oral language (standard scores), phonological processing (standard scores), and rapid automatized naming (scale scores) skills measured at school-age. **B.** The association results between school-age skill performance and each of the infant FCFs of the four identified seed regions, including the left temporal pole (the middle portion, midLTPO), left fusiform gyrus (LFFG), right fusiform gyrus (RFFG) and right inferior temporal gyrus (RITG). The distribution maps for each pair show the performances of the true prediction analyses (red) against the null distribution (blue) based on permutation results. The identified longitudinal associations were further revaluate using a leave-one-out cross-validation approach where one subject’s dataset was selected for testing while the remaining subjects were applied to train the support vector regression model. The obtained SVR results are presented as a scatter plot in the upper right corner of each cell to illustrate the fitness of true performance and predicted outcomes across the whole sample (see detailed analysis procedure and the statistical results in SI Methods and Table S5). Significant associations (Bonferroni corrected) with a large effect (Cohen’s d ≥ 0.8) and a 95% bootstra confidence interval above zeros were highlighted in red.

### Early functional network correlates of long-term verbal development are established within the first year of life

Our first analyses revealed strong prospective associations of school-age performance in the FC patterns of infants 4 and 18 months old. Since functional neural networks develop rapidly during the first two years of life [63], the age range of the imaging collection time point might lead to the assumption that the observed relationship could be driven by the more established functional networks among the older infants (>12 months old). Therefore, to further evaluate the importance of the early-emerged functional connectivity patterns in the observed associations, we reran the SVR analyses in the four identified FCFs using a subset of 29 infants who completed the imaging session before one year of age (mean age = 8.6 ± 1.8 months). Based on the same criteria, strong associations were still present between the infant FCF of midLTP and oral language skills at school-age (*r* = 0.46 ± 0.37, Cohen’s *d* = 1.2, 95% BCI= [0.44, 0.49], *p*_corrected_ < 0.001) as well as those of left fusiform gyri (*r* = 0.39 ± 0.42, Cohen’s *d* = 0.88, 95% BCI= [0.37, 0.42], *p*_corrected_ < 0.001) and RITG (*r* = 0.33 ± 0.43, Cohen’s *d* = 0.82, 95% BCI= [0.31, 0.36], *p*_corrected_ < 0.001) for the phonological skills measured at age 6. However, the association between the infant FCF of RFFG and the subsequent phonological skills (*r* = 0.36 ± 0.43, Cohen’s *d* = 0.74, 95% BCI= [0.33, 0.39], *p*_corrected_ < 0.001) failed to outperform permutation results by large effect (Cohen’s *d* > 0.8) and therefore was not considered in the subsequent analyses.

### Prospective associations between infant FCFs and school-age language and phonological performance are independent of early language milestones

Foundational language skills have been demonstrated to emerge early in infancy, which lay a critical foundation for (and are often associated with) the development of more complex language functions (see [64] for a review). It is, therefore, critical to address the question as to whether the emergent language skills of these infants might explain the observed longitudinal links between the FCFs in infancy and school-age performance. To address this, we further selected a subset of 24 infants from the original sample, whose receptive and expressive language skills were assessed at the infant time point. We then examined whether the inclusion of language skills measured in infancy change the prospective associations between infant functional networks and school-age performance identified in previous analyses; namely, the infant FCF of midLTP and oral language skills, and the infant FCFs of LLF and ITG and phonological skills. Specifically, two SVR models were run for each of the three identified associations. While both models included regular covariates similar to the ones included in the main analyses, only the second model further considered the receptive and expressive language skills measured in infancy. The association distributions derived from the two models were compared using the Kolmogorov-Smirnov test. If the identified prospective associations are *partially* mediated by language skills in infancy, we should expect significant decreases in association performances of the model controlling for infant language skills compared to the model that doesn’t control for infant language skills.

Results showed no reduction in the association strength between the identified FCFs and school-age oral language (midLTP: *p_uncorrected_* = 0.21) and phonological skills (LFFG: *p_uncorrected_* = 0.28; RITG: *p_uncorrected_* = 0.93) when the infant language skills were included as covariates compared to without (Figure 3). Lastly, correlation analyses were also run between each of the school-age language and pre-literacy scores and their infancy MSEL scores for the receptive and expressive language skills, which did not reveal any significant associations, even before correction for multiple comparisons (all *p*_uncorrected_ > 0.1, Table S6).

**Figure 3.**
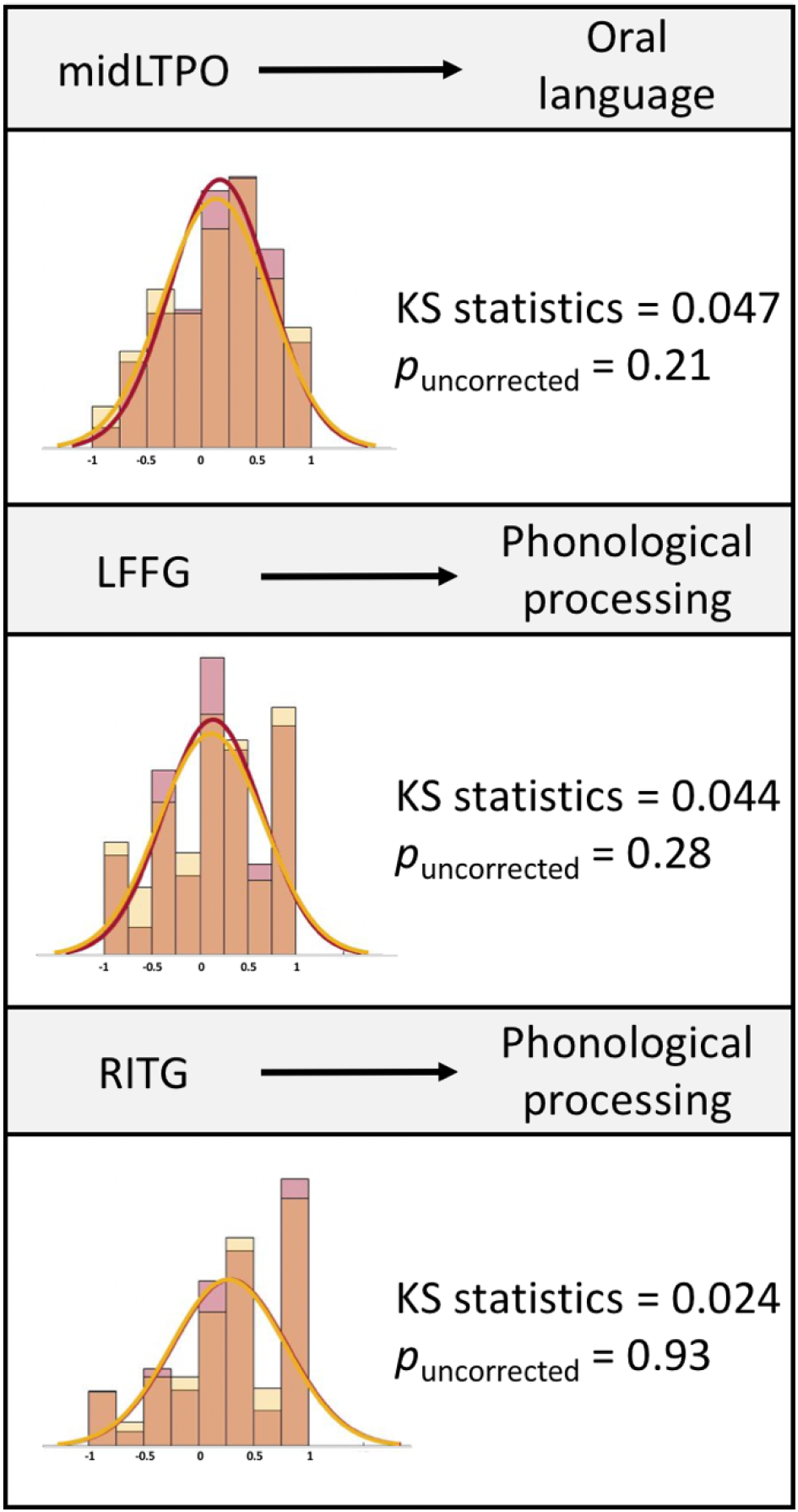
No significant differences in SVR performance with (yellow) and without (red) infant language milestones included as control variables. To evaluate whether (pre)language skills in infancy might contribute to the identified prospective associations, we further compared two models with and without receptive and expressive language skills in infancy included as covariates (in addition to the other measures included in the previous analyses) respectively. A Kolmogorov Smirnov (KS) test was used to compare th distributions of association strengt between the two models. No significant results were obtained before correcting for multiple comparisons. Abbreviations: midTPO = Temporal pole (the middle portion); LFFG = Left fusiform gyrus; RITG = Right inferior temporal gyrus.

## Discussion

In this study, we provide the first evidence for prospective associations between functional network characteristics in infancy and school-age development of language and foundational literacy skills. Utilizing a five-year longitudinal imaging dataset, we identify three temporal regions whose functional connectivity fingerprint patterns within the first year of life could reliably account for individual differences in oral language (left temporal pole) or phonological skills (left fusiform gyrus and right inferior temporal gyrus) measured at school-age in the same group of children. These strong associations are observed when the critical environmental factors of SES and HLE in infancy, as well as general cognitive abilities at school-age, were controlled, suggesting the early emergence of neural networks for long-term language and phonological development. We further show that these longitudinal links are independent of (pre)language skills in infancy, suggesting that the neural mechanisms underlying subsequently developing language and pre-literacy skills might be, at least partially, rooted early in the emergent functional network architecture of infant brains. Altogether, these findings uncover a critical role of functional connectivity patterns established in infancy on the protracted development of high-order cognitive functions of language and literacy.

The current study utilizes functional connectivity fingerprints to identify brain regions associated with subsequent cognitive development, highlighting the importance of inter-regional interactions in regional functional specialization through development. It has been long hypothesized that the functional development of individual regions is facilitated and shaped by earlier developing connectivity patterns [57, 65]. In line with this account, it has been shown that the functional connectivity pattern of one region at rest is associated with task-based fMRI activation of this region [66], and changes in the FC patterns parallel the anatomical boundaries between functionally distinct neighbors [67]. Moreover, it has been recently demonstrated that the (structural) connectivity fingerprints at age five before children learned to read are predictive of word-specific functional activation after 3-years’ reading instruction [68], providing direct evidence for the developmental significance of network mechanisms for neural functional specialization. Following the same line, our current study demonstrates that functional connectivity pattern established in infancy for a given region is associated with the cognitive skill this region subsequently supports (see more discussion below), suggesting that functional topologies that emerge within the first year of life lay a critical neural foundation for subsequent long-term development. However, future studies are needed to further examine the role of early-developed functional connectivity in shaping the neural correlates of cognitive functions.

The infant FCF pattern of the left temporal pole is associated with the school-age language performance, shedding light on an early functional network mechanism that could underlie the acquisition of complex language skills. Previous fMRI studies have revealed neural activation in the left temporal pole during speech perception in infants (e.g., [38, 39, 60]). In adults, the temporal pole is involved in word- and picture-based semantic tasks (e.g., [69–72]), and typically viewed as a hub for lexico-semantic representation [73, 74]. Therefore, our working hypothesis is that the temporal pole might serve as a core component of a wide-spread neural substrate that enables the learning of lexico-semantic representation via early language and speech learning/processing. Moreover, the fact that this association is observed when key environmental factors and the school-age general cognitive skills are controlled for and is also independent of (pre)language abilities in infancy further highlights that the early-developed neural networks underlie high-order language acquisition. Our hypothesis is also consistent with Dehaene-Lambertz and colleagues’ [75] recent proposal of an early symbolic system associated with speech/language learning observed in infants from the first months of life onward. In summary, the current finding provides empirical evidence of the far-reaching developmental significance of this early-developing functional network on the trajectory of language acquisition. Future work is warranted to further investigate the network characteristics and specific roles of their constituent brain regions in supporting this complex learning process.

We further identify prospective associations between pre-literacy skill, i.e., phonological processing and the infant FCF patterns of the left fusiform gyrus and right inferior temporal cortex, regions relevant for literacy and language processing. The left fusiform gyrus is central to literacy development. It hosts the visual word form area, which is developed after the onset of learning to read and underlies the orthographic processing of words [76]. Moreover, the fusiform gyrus is also activated during phonological processing (e.g., [77]) and its neural activation pattern encodes both phonological and orthographic information of words [78]. Here, it is demonstrated that functional connectivity patterns of the left fusiform gyrus established with the first year of life are prospectively associated with the development of phonological skills in school-age children, suggesting a functional role in the development of phonological and orthographic processing. Moreover, the bilateral inferior temporal gyri (and fusiform to a lesser extent) are also important components of the ventral stream within the neuroanatomy of speech processing, which serve as an interface linking speech code (i.e., phonological representation) with the corresponding semantic information [79, 80]. Brain lesion sizes to this area (inferior temporal and fusiform gyri) have been shown to associate with the auditory word comprehension performance among stroke patients, even after controlling for the conceptual knowledge as indexed by performance in object recognition [81]. Greater activation of this area has been observed when participants listened to semantically-ambiguous sentences compared to unambiguous sentences [82, 83]. Therefore, the observed relationship between the functional network of the ventral speech processing pathway and phonological development is consistent with the longitudinal associations between the development of speech perception and oral language comprehension at infant and toddler stages and literacy skills at school-age [44, 84–86]. Lastly, it is also interesting to note that rapid automatized naming (RAN) abilities measured at school-age failed to be reliably associated with FCF of any temporal region. The RAN assessment measures the efficiency of integrating multiple language-related (e.g., access and retrieval of phonological and semantic information) and general cognitive (e.g., attention, vision, and motor planning) processes [54]. Therefore, the null results might indicate that the RAN skills reflecting the integration abilities might be minimally associated with early-emerged functional networks in infancy.

Broadly speaking, our study illustrates the importance of combining developmental psychological knowledge with infant imaging techniques to characterize the initial “blueprints” of the brain’s functional organization that support the protracted development of complex cognitive function. High-order cognitive functions, such as language and reading, are complex systems that involve orchestration of multiple distinct yet interactive cognitive processes (e.g., [87]). These components are supported by specific neural mechanisms in adults (e.g., [88, 89]), and *often* exhibit characteristic neural functional specialization trajectories. For example, while neural responses of speech perception are evident in newborns (e.g., [38]), functional tuning of visual letter recognition is only demonstrated after the onset of reading instruction [90]. Therefore, surveying the relationship between infant brain characteristics and the subsequent development of high-order cognitive abilities within a cognitive framework enables the depiction of structured cortical scaffolding, upon which cognitive abilities emerge. In line with this, our current study extends the previous observation of the associations between infant functional network and overall language development at age 4 [51] by demonstrating cognitive process-specific relationships between functional connectivity fingerprints in infancy and school-age performance of oral language and literacy skills. Overall, such an interdisciplinary approach integrating developmental psychology, cognitive sciences and pediatric (infant) imaging methodologies can provide a valuable opportunity to identify early brain functional characteristics linked to individual variation in core processes of high-order cognitive functions, highlighting the potential of developing imaging markers to identify specific weakness at-risk populations for targeted early intervention [63].

Our study represents the first evidence of prospective associations between infant functional topologies and long-term development of language and foundational literacy skills measured at school-age. However, the results should be interpreted with some caution. The infants investigated in the current study had a wide age range at our initial scan. Due to challenges associated with the recruitment and retention of a longitudinal infant MRI cohort, participants with a wide range of age (4-18 months old) at the initial scan were included. This imposes challenges for both data processing and result interpretation. Based on the mean age of the infant sample (10.2 ± 3.5 months), the one-year-old AAL atlas was utilized in the current study [58]. Due to the significant growth and myelination of the human brain within the first two years of life (e.g., [91–93], applying one atlas to an infant sample of 4-18 months old could potentially introduce registration errors. To minimize this risk, visual inspection was conducted to ensure the correct mapping of every cortical region to the structural image of each infant. Moreover, despite the risk of overlooking functional subdivisions within each cortical region, the current approach of adopting the mean time series of cortical regions defined by anatomical parcellation is more resistant to registration errors than a more fine-grained parcellation derived from a different age group. Ultimately, the development of age- /developmental stage-specific atlases is needed and will be crucial to better characterize and understand the functional network mechanisms underlying cognitive development in infants (see [94, 95] for reviews). Furthermore, several approaches were adopted to minimize the potential confounding effects of age-at-scan, including modeling age-at-scan and the infant’s language environments as covariates in the main analyses, performing conformational analyses in a subgroup of younger infants, and directly evaluating the contribution of early foundational language skills on observed associations. Nevertheless, the current project design limits a clear distinction between neural mechanisms established at birth and contributions of postnatal growth/experiences. Future longitudinal investigation starting in neonates, along with behavioral and neural characterization at multiple developmental stages in a longitudinal design, are needed to unravel neural trajectories underlying language and literacy development and key components that shape this process.

## Conclusion

Utilizing a five-year longitudinal neuroimaging project starting in infancy, the current study provides the first empirical evidence of prospective associations between functional connectivity fingerprints in infants and the development of oral language and phonological skills in school-age children. Moreover, we further show that these longitudinal associations with school-age language and pre-literacy skills are not explained by foundational language skills in infancy, suggesting a unique role of functional networks in the development of language and foundational literacy abilities. Overall, this body of work reveals early-emerged functional network mechanisms in infancy that underlie the long-term development of complex cognitive abilities with a protracted neuro-trajectory that starts early in life or even in utero.

## Methods

### Participants and study overview

All infants were recruited at Boston Children’s Hospital (BCH) as part of an ongoing longitudinal project which aims to characterize brain changes underlying language and reading development from infancy to school-age. The current protocol was reviewed and approved by the institutional review board at BCH. Before participation, informed written consent was obtained from a parent or legal guardian of each infant.

A total of 60 infants (4-18 months) successfully completed resting-state functional and structural MR imaging between the years of 2011 and 2015. Among them, six subjects were excluded from further analysis due to poor image quality (n = 2) or excessive motion (n = 4, see details below). Among 54 subjects with usable imaging data, 42 (21 females, 10.2 ± 3.5 months) were successfully followed over five years (5.6 ± 1.0 years) until school-age (mean age = 6.5 ± 0.96 years). Infant participants who were successfully followed (n=42) and those were not (n=12) did not differ significantly by age, gender, head movement during scanning, emergent language skills in infancy, or environmental measures (all *p_u_*_ncorrected_ > 0.05, table S6). The longitudinal sample, characteristically representing the whole sample, was thus included in the current prospective association analyses. All 42 children were from English-speaking households and were exposed to the English language since birth. They further demonstrated typical general cognitive ability based on their performance (standard scores (SS) > 85, mean SS = 107 ± 13) on the matrices subtest of the Kaufman Brief Intelligence Test-II at school-age (KBIT-II, Kaufman & Kaufman, 2004. Moreover, none of the subjects had birth complications, neurological trauma, or reported developmental delay. All subjects had MRIs that were clinically interpreted as normal by a pediatric neuroradiologist at BCH.

#### Psychometric and environmental measures

At the infant stage, emergent receptive and expressive languages were evaluated using the Mullen Scales of Early Learning assessment (MSEL, [96]) in 29 infants. Raw scores were converted into T scores with a mean of 50 and SD of 10 for the subsequent analyses. Furthermore, environmental factors critical for language and literacy development were also collected through parental questionnaires, which included socio-economic status (SES, i.e., parental education, n = 41) and home literacy environment (HLE, adapted from [97], n = 38, see SI Methods).

At the school-age stage, a comprehensive assessment battery was administered to evaluate each child’s general cognitive ability, language, literacy, reading, and spelling skills. Since most of the child participants had not yet developed reading and spelling skills during the second longitudinal timepoint (mean age = 6.5 ± 0.96 years), the present study focused on language and foundational literacy skills that could be reliably measured within this age range. This included assessments of oral language (OL, the OL subtest of the Woodcock-Johnson-IV assessment (WJ-IV), [98]), rapid automatized naming (RAN, Comprehensive Test of Phonological Processing-II (CTOPP-2), [99]) and phonological processing skills (the phonological processing subtest of WJ-IV, see detailed descriptions of assessments in the SI Methods).

#### Image acquisition

All images were acquired during natural sleep without sedation on a Siemens 3T Trio MRI scanner with a 32-channel adult head coil (see [100] for a detailed infant imaging procedure). A motion-compensated T1 weighted multi-echo MPRAGE sequence was applied to collect high-quality structural images for each infant (slice number = 128, TR = 2520 ms, flip angle = 7°, field of view = 192 mm^2^, voxel size =1 × 1 × 1 mm^3^). Moreover, an 8-minute EPI-BOLD sequence was applied to collect resting-state fMRI data for every infant (TR = 3000 ms, TE = 30 ms, flip angle = 60°, voxel size =3 × 3 × 3 mm^3^).

#### Image analysis

##### a. Resting-state fMRI preprocessing

Functional MRI images were first corrected for slice timing using FSL/SLICETIMER (and an in-house function for images with multi-slice acquisition). Rigid body alignment for motion correction and estimation of motion regressors was performed using FSL/MCFLIRT. Anatomical and functional MRI volumes were spatially normalized to the UNC 1-year old infant template [58] using affine transformations (FSL/FLIRT). After spatial normalization, linear regression was performed using the following nuisance regressors: (1) six motion regressors from realignment estimates (accounting for translations and rotations in three directions), (2) mean cerebral spinal fluid (CSF) and white matter (WM) signals extracted from subject-specific anatomical masks (computed with FSL/FAST), and (3) spikes regressors based on motion outliers (see §Artifact detection). Temporal band-pass filtering (0.01-0.1Hz) and spatial smoothing (Gaussian filter, FWHM = 6 mm) were applied after nuisance regression.

#### Artifact detection

An in-house script was used to identify artifacts in the fMRI data, including motion and spiking (https://github.com/xiyu-bnu/infant_restingstate_prediction). Framewise displacement (FD) was estimated from translations and rotations obtained from rigid-body alignment after motion correction [101]. Volumes with FD > 0.3 mm were considered outliers. In order to minimize potential artifacts in neighboring volumes due to temporal filtering or spin history, one preceding and two subsequent frames were also marked as outliers. A binary vector indexing all the outliers was built for each subject and used as a motion regressor to remove contaminated frames from the time series prior to filtering. Only subjects with more than 5-minute usable volumes were considered for further analysis.

##### b. Seed-based connectivity analysis

Seed regions were based on 78 cortical regions of interest (ROIs) defined by the 1-year-old Infant Brain Atlases [58] derived from the Automated Anatomical Labeling (AAL) atlas [102]. Seed-based time series were extracted by averaging the BOLD signal across all the voxels in each ROI. The functional connectivity fingerprint (FCF) pattern associated with each seed ROI was generated by correlating the BOLD time series of the ROI with that of all other ROIs using the Pearson product-moment correlation formula. Correlation coefficients were normalized to z-scores using Fisher’s transformation.

##### c. Prediction

A Support Vector Regression (SVR) classifier was applied for the prospective association analyses using the LIBSVM package [103]. Relationships between the FCF pattern of each anatomical region in the temporal lobe and the school-age performance of each assessment (i.e., OL, phonological processing, and RAN) were examined. A cross-validation approach was applied in the SVR analyses, in which 20% of the datasets were held out (testing dataset) while the remaining datasets were used to train an SVR model. Moreover, to minimize potential confounding effects on the association analysis, a linear regression model was first fit onto the training dataset with task performance (standardized scores) as the dependent variable, and environmental factors (parental education level and HLE), age at the infant scan, and general cognitive ability at school-age as independent variables. The residual values of each regression were then used as outcome measures in the SVR analyses. The obtained regression model was further applied to the testing dataset in order to obtain the residual values corresponding to the task performance for the testing participants. The trained SVR model was then applied to the testing sample to produce the predicted values, which were further correlated (Spearman) with the real performance (residual values) as the SVR performance. This process was repeated 1000 times to generate the true distributions of the SVR performance. The distribution of the prediction performance was tested against a null or random distribution, which was generated following the same procedure except that the test performance used during training was randomly permuted in each iteration. Statistical significance against the null distribution was tested with a Kolmogorov-Smirnov test. The effect size between the true and null distributions was also measured using Cohen’s *d* measure [62], where larger magnitudes indicate higher predictive power of the infant FCFs on school-age performance when compared to the null distribution. In addition, a 95% bootstrap confidence interval (BCI) for the mean association strength was computed based on the true distribution. Only associations significantly (corrected for the comparison comparisons) outperforming the null distribution with a large effect size (Cohen’s *d* > 0.8, [62] and a 95% BCI values above zero were considered in the current analyses (see Figure 1 for a general pipeline of the prediction analyses).

## Data availability

All documentation and code used in these analyses can be shared with any research team. Due to Institutional Review Board regulations at Boston Children’s Hospital at the time of consent, our data cannot presently be uploaded to a permanent third-party archive. However, data sharing can be initiated through a Data Usage Agreement. We are strongly committed to working closely on such a sharing plan with anyone who would like to re-analyze our data and will provide templates and procedural outlines.

## Acknowledgments

This study was supported by the Eunice Kennedy Shriver National Institute of Child Health and Human Development #R01HD65762-01 (awarded to N.G. and P.E.G), the Charles H. Hood Foundation (awarded to N.G.), and the Boston Children’s Hospital Pilot Grant (awarded to N.G.).

We sincerely thank all the families for their participation in this longitudinal study.

## Competing interests

The authors declare no competing financial interests.

## Supporting Information

### SI Methods

#### Characterization of familial environment at the infant time point

The environmental characteristics of participating families, including socio-economic status (SES) and home literacy environment (HLE), were collected through the parental questionnaire completed during the first visit at the infant time point.

The SES was characterized in terms of the education background of both parents, which were regarded as a more stable indicator of SES compared to other frequently used variables, such as parental income and occupation (Bradley & Corwyn, 2002; Saifi & Mehmood, 2011). Both parents’ highest education degrees were recorded on a six-point scale (see table S1) and averaged as a composite score of SES for each infant. Regarding the HLE measures, the responses for each of the ten questions were first converted into z-scores, and then averaged as one composite score of HLE per infant (see table S1 for the distribution of answers for each of the SES- and HLE-related questions).

#### Evaluation of language and preliteracy skills at school-age

Non-verbal general cognitive ability was assessed using the Matrices subtest of KBIT-II, which evaluates children’s fluid reasoning skills using picture-based stimuli with minimal requirement of verbal knowledge. Participants’ oral language skills were examined using the Oral Language Cluster of the WJ-IV Test of Oral Language. This cluster is comprised of two subtests: 1) the Picture Vocabulary subtest in which participants are asked to identify pictured objects, and 2) the Oral Comprehension subtest in which children are asked to listen to oral passages and fill in final missing words based on the semantic and syntactic information provided. Moreover, each participant was evaluated on their RAN and phonological processing skills, two foundational literacy skills that are among the most commonly identified cognitive precursors of reading outcomes. RAN refers to the ability to name a series of stimuli as quickly and accurately as possible and has been linked to automaticity of retrieval and subsequent reading fluency (Lervåg & Hulme, 2009; Norton & Wolf, 2012). Every child received three RAN subtests from the CTOPP-2 assessment, each using pictured objects, colors, and letters, respectively. Phonological skills were examined using three subtests included in the Phonological Processing Cluster of the WJ-IV Test of Cognitive Abilities. They were 1) Word Access, which required participants to provide a word that had a particular sound in the beginning, middle, or end; 2) Word Fluency, which examined the ability to produce the maximal number of words beginning with a certain letter within one minute; and 3) Substitution, in which the child must substitute sounds in words to make new words (say “can” but with a “t” instead of an “n”). Raw scores acquired for each task were converted into standard scores for subsequent analyses. For the oral language and phonological processing clusters of the WJ-IV Test that were evaluated using multiple subtests, one composite score was calculated for each cognitive cluster following the guidelines in the assessment manual. Moreover, an average RAN score was obtained for each subject by averaging across performance of all included subtests (RAN).

#### The leave-one-out cross-validation (LOOCV) approach

The Support Vector Regression (SVR) analyses with LOOCV followed the same procedure as described in the main manuscript, except that during each iteration, one subject, instead of a group of subjects (20% in our case) was used as testing sample. Accordingly, all the remaining subjects (n-1) datasets were applied to train the SVR model. The same procedure repeated until all subjects were used for testing once. Correlation analyses were performed between the predicted and real performance across the whole sample, and the corresponding correlation coefficient was considered as the SVR performance. Moreover, to evaluate the reliability of the SVR performance, permutation tests were further carried out with the same analysis steps, but the task performance was randomly assigned to each subject. These analyses repeated 1000 times, generating a null distribution. The significance of the SVR model was thus determined as the percentage of instances in which the correlation coefficients based on the permutation tests were larger or equal to the one derived from the SVR model based on the real outcome measures. Results obtained here (see Table S4) were similar to the findings reported in the main manuscript based on the validation approach using separate testing and training samples.

---

**Table S1.**
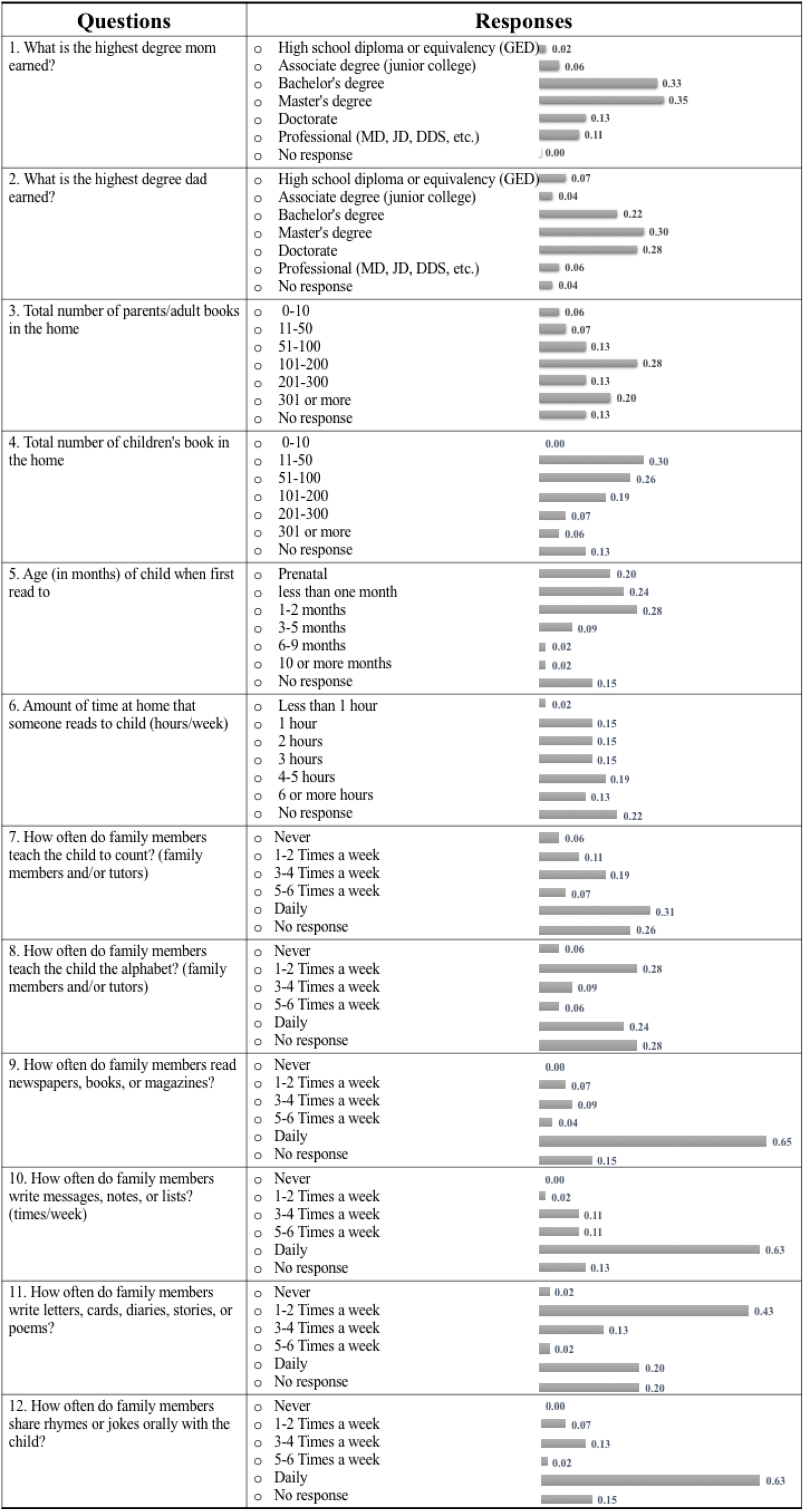
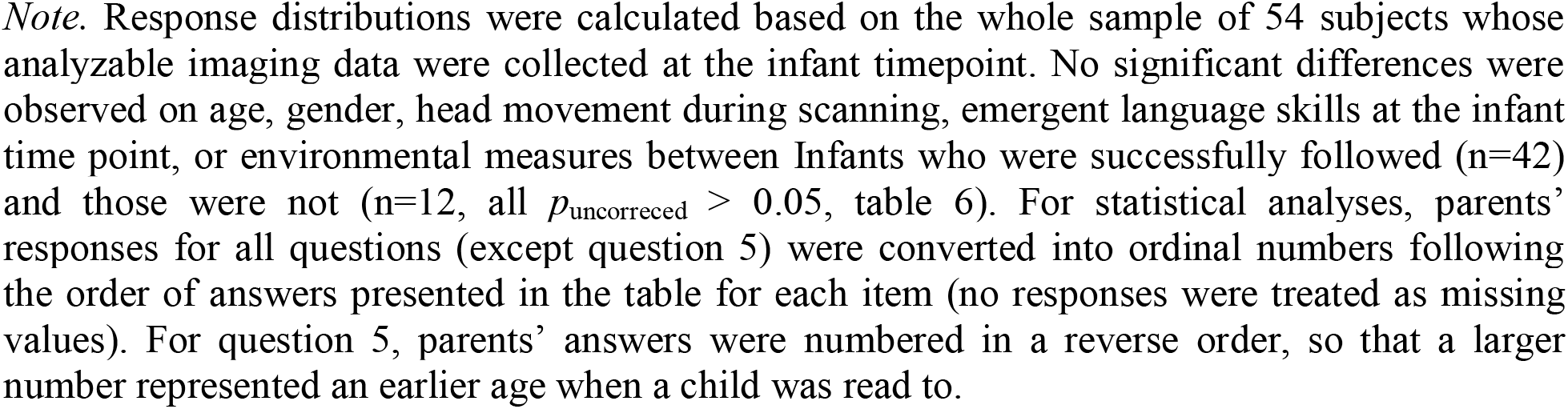
Parental responses for the survey questions on the socio-economic status and home literacy environment for infant participants

| Questions | Responses |
| --- | --- |
| 1. What is the highest degree mom earned? | High school diploma or equivalency (GED) 0.02<br>Associate degree (junior college) 0.06<br>Bachelor's degree 0.33<br>Master's degree 0.35<br>Doctorate 0.13<br>Professional (MD, JD, DDS, etc.) 0.11<br>No response 0.00 |
| 2. What is the highest degree dad earned? | High school diploma or equivalency (GED) 0.07<br>Associate degree (junior college) 0.04<br>Bachelor's degree 0.22<br>Master's degree 0.30<br>Doctorate 0.28<br>Professional (MD, JD, DDS, etc.) 0.06<br>No response 0.04 |
| 3. Total number of parents/adult books in the home | 0-10 0.06<br>11-50 0.07<br>51-100 0.13<br>101-200 0.28<br>201-300 0.13<br>301 or more 0.20<br>No response 0.13 |
| 4. Total number of children's book in the home | 0-10 0.00<br>11-50 0.30<br>51-100 0.26<br>101-200 0.19<br>201-300 0.07<br>301 or more 0.06<br>No response 0.13 |
| 5. Age (in months) of child when first read to | Prenatal 0.20<br>less than one month 0.24<br>1-2 months 0.28<br>3-5 months 0.09<br>6-9 months 0.02<br>10 or more months 0.02<br>No response 0.15 |
| 6. Amount of time at home that someone reads to child (hours/week) | Less than 1 hour 0.02<br>1 hour 0.15<br>2 hours 0.15<br>3 hours 0.15<br>4-5 hours 0.19<br>6 or more hours 0.13<br>No response 0.22 |
| 7. How often do family members teach the child to count? (family members and/or tutors) | Never 0.06<br>1-2 Times a week 0.11<br>3-4 Times a week 0.19<br>5-6 Times a week 0.07<br>Daily 0.31<br>No response 0.26 |
| 8. How often do family members teach the child the alphabet? (family members and/or tutors) | Never 0.06<br>1-2 Times a week 0.28<br>3-4 Times a week 0.09<br>5-6 Times a week 0.06<br>Daily 0.24<br>No response 0.28 |
| 9. How often do family members read newspapers, books, or magazines? | Never 0.00<br>1-2 Times a week 0.07<br>3-4 Times a week 0.09<br>5-6 Times a week 0.04<br>Daily 0.65<br>No response 0.15 |
| 10. How often do family members write messages, notes, or lists? (times/week) | Never 0.00<br>1-2 Times a week 0.02<br>3-4 Times a week 0.11<br>5-6 Times a week 0.11<br>Daily 0.63<br>No response 0.13 |
| 11. How often do family members write letters, cards, diaries, stories, or poems? | Never 0.02<br>1-2 Times a week 0.43<br>3-4 Times a week 0.13<br>5-6 Times a week 0.02<br>Daily 0.20<br>No response 0.20 |
| 12. How often do family members share rhymes or jokes orally with the child? | Never 0.00<br>1-2 Times a week 0.07<br>3-4 Times a week 0.13<br>5-6 Times a week 0.02<br>Daily 0.63<br>No response 0.15 |

*Note.* Response distributions were calculated based on the whole sample of 54 subjects whose analyzable imaging data were collected at the infant timepoint. No significant differences were observed on age, gender, head movement during scanning, emergent language skills at the infant time point, or environmental measures between Infants who were successfully followed (n=42) and those who were not (n=12, all $p_{\text{uncorrected}} > 0.05$ , table 6). For statistical analyses, parents' responses for all questions (except question 5) were converted into ordinal numbers following the order of answers presented in the table for each item (no responses were treated as missing values). For question 5, parents' answers were numbered in a reverse order, so that a larger number represented an earlier age when a child was read to.

**Table S2.**
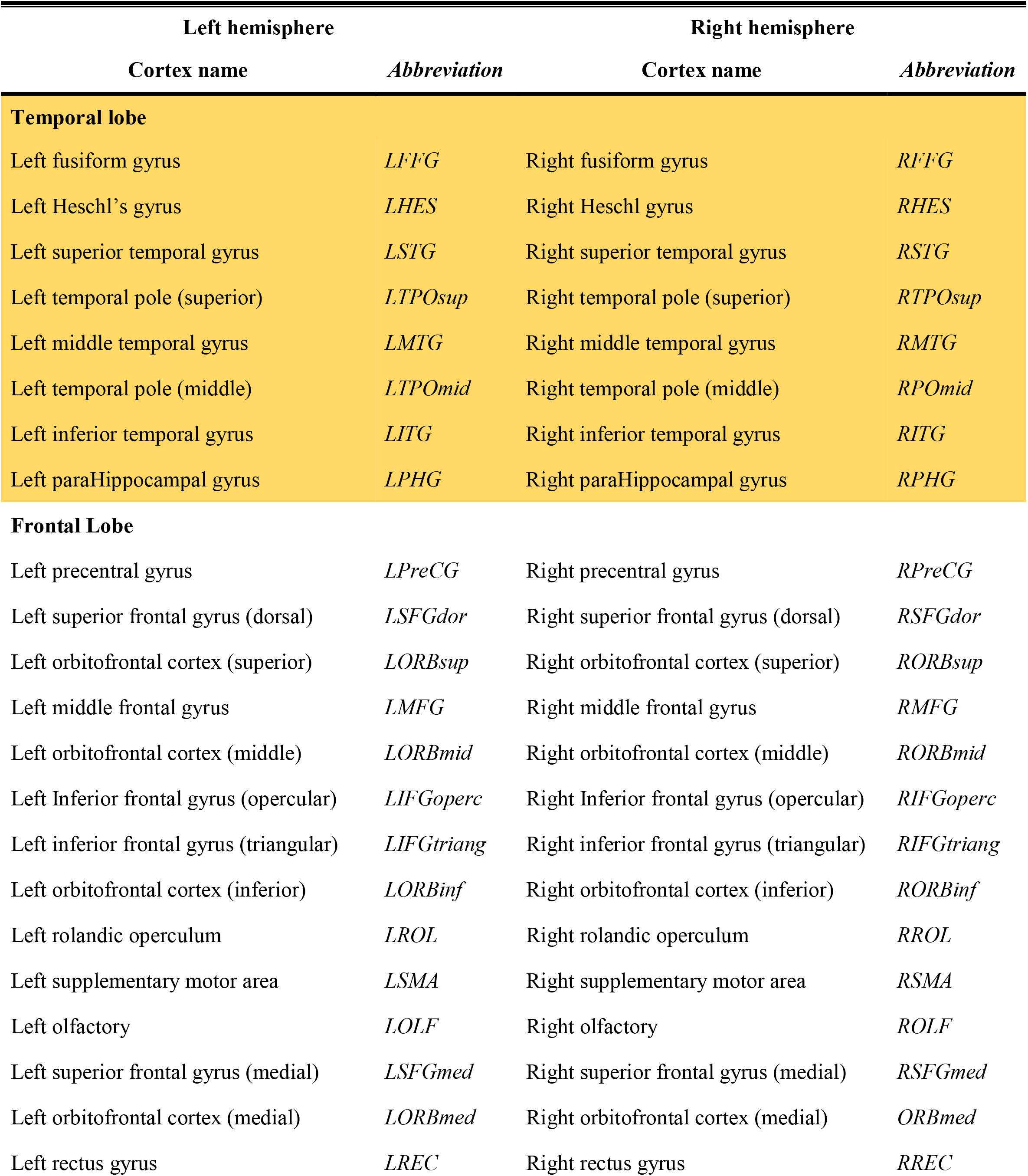
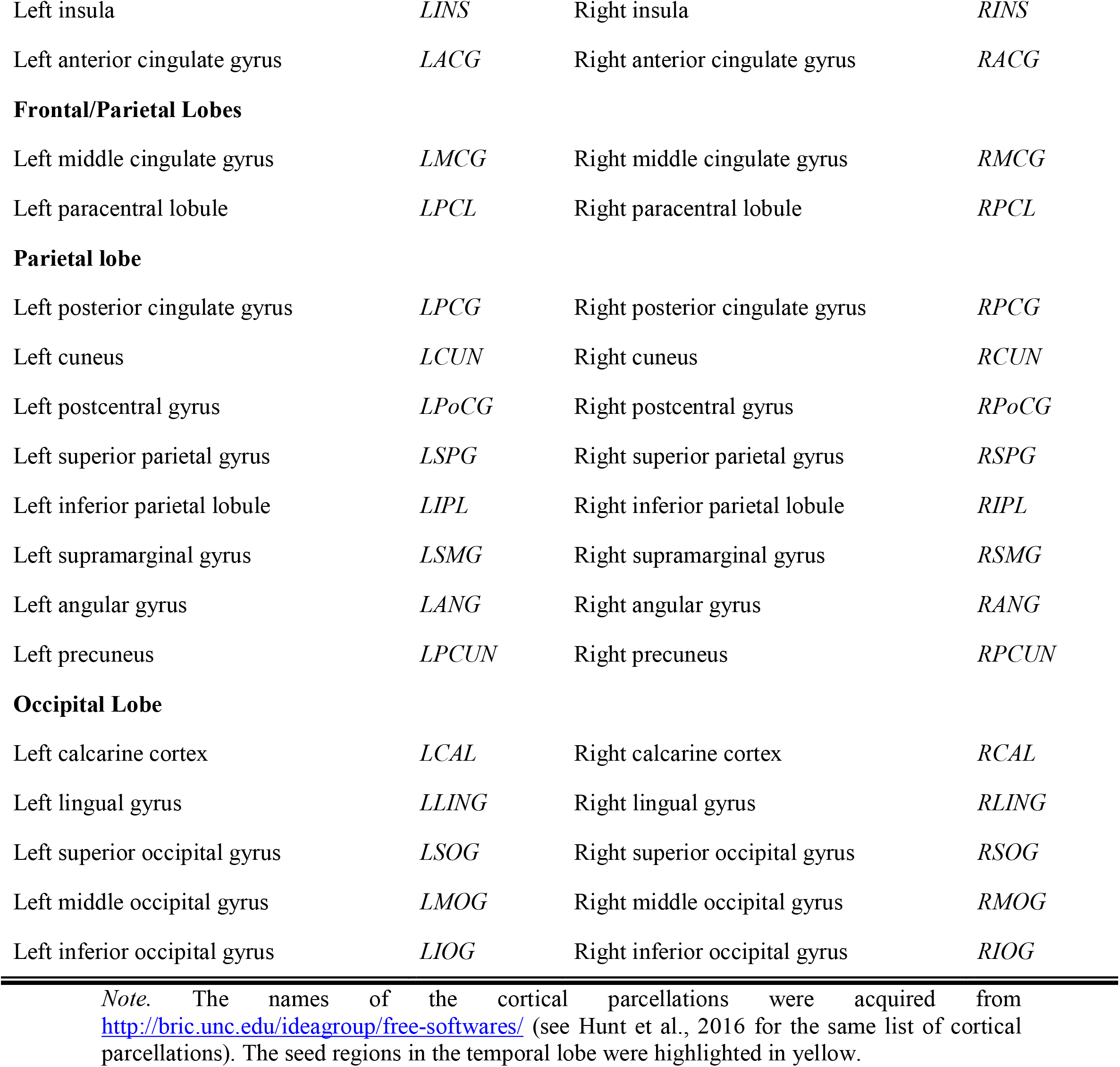
List of cortical parcellation derived from the 1-year-old infant AAL atlas (Shi et al., 2011)

**Table S3.** The full results of the SVR analyses based on the whole sample.

| Seed regions | Oral language |  |  | Phonological processing |  |  | Rapid automatized naming |  |  |
| --- | --- | --- | --- | --- | --- | --- | --- | --- | --- |
| | $r$ | Cohen's $d$ | 95% BCI | $r$ | Cohen's $d$ | 95% BCI | $r$ | Cohen's $d$ | 95% BCI |
| Right parahippocampal gyrus | $0.00 \pm 0.41$ | -0.04 | [-0.02, 0.03] | $0.01 \pm 0.39$ | -0.01 | [-0.01, 0.04] | $-0.31 \pm 0.31$ | -0.79 | [-0.33, -0.29] |
| Left parahippocampal gyrus | $0.09 \pm 0.38^*$ | 0.20 | [0.06, 0.11] | $0.17 \pm 0.39^*$ | 0.36 | [0.15, 0.19] | $-0.18 \pm 0.37$ | -0.45 | [-0.20, -0.16] |
| Right fusiform gyrus | $-0.15 \pm 0.36$ | -0.41 | [-0.17, -0.13] | $0.31 \pm 0.39^*$ | 0.82 | [0.29, 0.34] | $-0.22 \pm 0.34$ | -0.56 | [-0.24, -0.20] |
| Left fusiform gyrus | $0.15 \pm 0.40^*$ | 0.41 | [0.12, 0.17] | $0.40 \pm 0.37^*$ | 0.96 | [0.38, 0.43] | $-0.05 \pm 0.36$ | -0.16 | [-0.07, -0.03] |
| Right Heschl's gyrus | $0.19 \pm 0.38^*$ | 0.48 | [0.17, 0.21] | $0.05 \pm 0.42$ | 0.18 | [0.02, 0.08] | $0.00 \pm 0.40$ | 0.02 | [-0.02, 0.02] |
| Left Heschl's gyrus | $0.20 \pm 0.38^*$ | 0.47 | [0.18, 0.22] | $-0.01 \pm 0.42$ | -0.01 | [-0.04, 0.01] | $-0.08 \pm 0.37$ | -0.25 | [-0.10, -0.05] |
| Right superior temporal gyrus | $-0.39 \pm 0.31$ | -1.06 | [-0.41, -0.37] | $0.02 \pm 0.43$ | 0.08 | [-0.00, 0.05] | $0.10 \pm 0.37^*$ | 0.20 | [0.07, 0.12] |
| Left superior temporal gyrus | $-0.40 \pm 0.33$ | -1.15 | [-0.42, -0.38] | $-0.06 \pm 0.41$ | -0.19 | [-0.09, -0.04] | $-0.02 \pm 0.37$ | -0.09 | [-0.04, 0.00] |
| Right temporal pole (superior) | $0.19 \pm 0.36^*$ | 0.51 | [0.17, 0.22] | $-0.10 \pm 0.40$ | -0.20 | [-0.12, -0.07] | $-0.11 \pm 0.37$ | -0.30 | [-0.13, -0.08] |
| Left temporal pole (superior) | $0.02 \pm 0.38$ | 0.07 | [-0.0, 0.05] | $-0.15 \pm 0.44$ | -0.37 | [-0.18, -0.12] | $0.12 \pm 0.35^*$ | 0.28 | [0.10, 0.14] |
| Right middle temporal gyrus | $0.14 \pm 0.36^*$ | 0.32 | [0.12, 0.16] | $0.02 \pm 0.40$ | 0.04 | [-0.007, 0.04] | $0.01 \pm 0.36$ | 0.01 | [-0.01, 0.03] |
| Left middle temporal gyrus | $0.30 \pm 0.34^*$ | 0.71 | [0.27, 0.32] | $-0.11 \pm 0.39$ | -0.31 | [-0.14, -0.09] | $-0.10 \pm 0.36$ | -0.23 | [-0.12, -0.07] |
| Right temporal pole (middle) | $0.17 \pm 0.37^*$ | 0.47 | [0.15, 0.19] | $0.12 \pm 0.43^*$ | 0.28 | [0.09, 0.15] | $-0.08 \pm 0.38$ | -0.14 | [-0.10, -0.05] |
| Left temporal pole (middle) | $0.42 \pm 0.36^*$ | 1.09 | [0.40, 0.44] | $0.06 \pm 0.41$ | 0.12 | [0.038, 0.09] | $0.05 \pm 0.37$ | 0.09 | [0.02, 0.07] |
| Right inferior temporal gyrus | $0.04 \pm 0.36$ | 0.06 | [0.01, 0.06] | $0.41 \pm 0.36^*$ | 0.94 | [0.39, 0.43] | $-0.03 \pm 0.38$ | -0.07 | [-0.06, -0.01] |
| Left inferior temporal gyrus | $-0.10 \pm 0.35$ | -0.31 | [-0.12, -0.08] | $0.10 \pm 0.40^*$ | 0.25 | [0.08, 0.13] | $0.00 \pm 0.39$ | -0.04 | [-0.03, 0.02] |

*Note.* Results highlighted in red are reported in the manuscript, as they showed a large effect (Cohen's $d > 0.8$ ), exhibited a 95% BCI above zero, as well as significantly outperformed the respective randomized distribution.
\* SVR performance based on the real task performance significantly (after correcting for multiple comparisons) outperformed the null results derived from the randomized labels ( $p_{\text{corrected}} < 0.05$ ).

**Table S4.** SVR performance between identified functional connectivity fingerprints (FCF) and school-age performance using different proportions of training and testing datasets.

| FCFs in infancy | School-age performance | $r$ | Cohen's $d$ | 95% BCI |
| --- | --- | --- | --- | --- |
| SVR results derived from 70% training datasets and 30% testing datasets |  |  |  |  |
| Left temporal pole (middle) | Oral language | $0.42 \pm 0.24$ | 1.5 | [0.41, 0.44] |
| Left fusiform gyrus | Phonological processing | $0.40 \pm 0.29$ | 1.2 | [0.38, 0.42] |
| Right fusiform gyrus | Phonological processing | $0.29 \pm 0.30$ | 0.91 | [0.27, 0.31] |
| Right inferior temporal gyrus | Phonological processing | $0.39 \pm 0.27$ | 1.3 | [0.38, 0.41] |
| SVR results derived from 60% training datasets and 40% testing datasets |  |  |  |  |
| Left temporal pole (middle) | Oral language | $0.38 \pm 0.19$ | 1.7 | [0.37, 0.39] |
| Left fusiform gyrus | Phonological processing | $0.40 \pm 0.24$ | 1.5 | [0.39, 0.42] |
| Right fusiform gyrus | Phonological processing | $0.29 \pm 0.24$ | 1.0 | [0.27, 0.30] |
| Right inferior temporal gyrus | Phonological processing | $0.40 \pm 0.22$ | 1.4 | [0.38, 0.41] |

Note. All SVR performance based on the real task performance significantly (after correcting for multiple comparisons) outperformed the null results derived from the randomized labels ( $p_{\text{corrected}} < 0.05$ ).

**Table S5.** Re-evaluation of longitudinal associations between identified functional connectivity fingerprints (FCF) and school-age performance using the leave-one-out cross-validation approach

| FCFs in infancy | School-age performance | *r* | *p* |
| --- | --- | --- | --- |
| Left temporal pole (middle) | Oral language | 0.50 | 0.006 |
| Left fusiform gyrus | Phonological processing | 0.52 | 0.014 |
| Right fusiform gyrus | Phonological processing | 0.35 | 0.065 |
| Right inferior temporal gyrus | Phonological processing | 0.50 | 0.015 |

**Table S6.** Correlation results between the infancy behavioral characteristics and school-age performance.

| Infancy performance |  | School-age performance |  |  |
| --- | --- | --- | --- | --- |
|  |  | Oral language | Phonological processing | Rapid automatized naming |
| MSEL | Receptive language | $r_{22} = 0.21$<br>$p = 0.34$ | $r_{18} = 0.31$<br>$p = 0.18$ | $r_{20} = 0.006$<br>$p = 0.98$ |
| | Expressive language | $r_{24} = -0.06$<br>$p = 0.77$ | $r_{19} = 0.11$<br>$p = 0.63$ | $r_{22} = -0.26$<br>$p = 0.21$ |
| Head movement during scanning | Framewise displacement | $r_{40} = -0.02$<br>$p = 0.90$ | $r_{34} = 0.024$<br>$p = 0.89$ | $r_{38} = -0.047$<br>$p = 0.78$ |
| | Number of removed scans | $r_{40} = -0.18$<br>$p = 0.25$ | $r_{34} = -0.11$<br>$p = 0.54$ | $r_{38} = -0.012$<br>$p = 0.94$ |

*Note.* The behavioral characteristics in infancy include the emergent language skills measured by MSEL and head movement during imaging acquisition. The uncorrected p-values were reported here. MSEL: Mullen Scales of Early Learning assessment

**Table S7.** Infant characteristics of participants who were longitudinally followed and those were not

|  | Longitudinally followed children | Non-followed children | Two-sample comparisons |
| --- | --- | --- | --- |
| Demographic information |  |  |  |
| Age (month) | 10.2 ± 3.5 | 9.6 ± 3.1 | $t_{52} = 0.51, p = 0.61$ |
| Gender (Female/Male) | 21/21 | 7/5 | $\chi^2 = 0.26, p = 0.61$ |
| Head movement during resting-state imaging |  |  |  |
| # of outlier images (%) | 5.2 ± 7.5 | 9.6 ± 14.7 | $t_{52} = 1.4, p = 0.16$ |
| Framewise displacement of the remaining scans (mm) | 0.067 ± 0.016 | 0.065 ± 0.029 | $t_{52} = 0.28, p = 0.78$ |
| Emergent language scores at the infant time point |  |  |  |
| Receptive language | 44.2 ± 8.3 | 46.3 ± 4.0 | $t_{25} = 0.43, p = 0.67$ |
| Expressive language | 46.9 ± 7.5 | 54.7 ± 6.1 | $t_{27} = 1.7, p = 0.10$ |
| Environmental characteristics |  |  |  |
| Socioeconomic status (parental education) | 3.7 ± 1.0 | 4.3 ± 0.89 | $t_{51} = 1.6, p = 0.11$ |
| Home literacy environment (z-score) | -0.014 ± 0.49 | 0.015 ± 0.32 | $t_{45} = 0.17, p = 0.87$ |

*Note.* Emergent language skills at the infant timepoint were assessed using the Mullen Scales of Early Learning assessment (Mullen 1995), and standard scores (T scores) were applied for between-group comparisons and result presentation. Participants' responses were missing for several measures, which were taken into consideration in the subsequent statistical analyses.

## Notes

### Competing Interest Statement

The authors have declared no competing interest.

